# Genomic and phenotypic comparison of *Saccharomyces cerevisiae* and *Saccharomyces boulardii*

**DOI:** 10.1101/2025.09.08.674931

**Authors:** Hannah Elena Duffey, Karl Alex Hedin, Hitesh P. Gelli, Troels Holger Vaaben, Morten Otto Alexander Sommer

## Abstract

*Saccharomyces boulardii* is a widely used probiotic yeast with clinical efficacy against certain gastrointestinal disorders. Although genomically related to *S. cerevisiae*, the extent to which *S. boulardii* harbors distinct probiotic-relevant traits remains incompletely defined, particularly across commercially distributed strains. Here, we performed comparative genomic, physiological, and functional analyses of five *S. boulardii* strains and three *S. cerevisiae* strains, including baker’s and laboratory variants. *S. boulardii* strains shared conserved genetic features and exhibited a conserved chromosomal inversion on chromosome XVI, lower copy numbers of CAZyme genes, and lineage-specific amino acid substitutions in central and tryptophan catabolism pathways—potentially underlying elevated production of immunomodulatory metabolites. *S. boulardii* strains also exhibited enhanced acid tolerance, elevated acetate and succinate production, and robust immunomodulatory activity, including suppression of IL-8 secretion and NF-κB, and consistent activation of the aryl hydrocarbon receptor (AhR) compared to *S. cerevisiae*. In contrast*, S. cerevisiae* strains displayed greater bile salt tolerance and faster growth under aerobic and anaerobic conditions at both 30°C and 37°C but lacked consistent anti-inflammatory effects or AhR agonism. Metabolic and immunological phenotypes varied with oxygen availability and strain background. Despite high genomic similarity, *S. cerevisiae* and *S. boulardii* exhibit distinct functional capacities relevant to probiotic efficacy. These findings help define species- and strain-specific features that inform the development and regulatory evaluation of next-generation yeast probiotics.

**Importance:** The yeast *Saccharomyces boulardii* is widely used as a probiotic to support human gut health, yet the reasons behind its beneficial effects remain unclear. This study compares *S. boulardii* with its close relative, *Saccharomyces cerevisiae*, which is commonly used in baking and research but does not show consistent health benefits. By examining multiple strains, we found that *S. boulardii* possesses unique features that may explain its ability to survive harsh gut conditions and influence the body’s immune responses. In contrast, *S. cerevisiae* grows faster and withstands bile better but lacks the same protective effects. These findings highlight how small genetic and physiological differences between related organisms can lead to distinct impacts on health. Understanding these differences provides a foundation for developing next-generation probiotics and for setting standards in their evaluation and use.

## Introduction

*Saccharomyces cerevisiae* is one of the most extensively characterised eukaryotic model organisms, foundational to studies in genetics, metabolism, and cellular regulation(1). Its well-annotated genome and robust genetic tractability have made it a cornerstone of molecular biology and functional genomics (1, 2). In parallel with its research utility, *S. cerevisiae* plays a central role in industrial biotechnology (3). It is widely used in baking, brewing, and winemaking, where its high fermentation efficiency, ethanol tolerance, and metabolic plasticity are exploited at commercial scale. More recently, it has become a favoured platform for synthetic biology and metabolic engineering, supporting the production of high-value compounds such as biofuels, bioplastics, pharmaceuticals, and nutraceuticals (4–6). Its GRAS (generally recognised as safe) status, scalability, and resilience under industrial conditions make it a versatile chassis for diverse biomanufacturing applications.

A closely related taxonomic variant, *Saccharomyces cerevisiae* var. *boulardii*, was the first commercially sold yeast strain used as a probiotic in clinical settings (7). Although classified within *S. cerevisiae*, *S. boulardii* exhibits distinct phenotypic, genomic, and functional characteristics (8–10). It is routinely administered for the prevention and treatment of gastrointestinal conditions such as antibiotic-associated diarrhoea, *Clostridioides difficile* infection, and inflammatory bowel disease (7, 11). A defining feature of *S. boulardii* is its ability to remain viable and metabolically active in hostile gastrointestinal environments, including exposure to low pH, bile salts, digestive enzymes, and elevated temperatures up to 37°C (8, 9, 12). These attributes have been shown to be absent in some of the more commonly used laboratory and industrial *S. cerevisiae* strains, which generally lack the ability to persist and function in the gut (8, 9).

The commercial use of *S. boulardii* is dominated by the CNCM I-745 strain, produced by Biocodex and validated in over 90 randomised clinical trials (13). However, multiple manufacturers market *S. boulardii* supplements using distinct, often proprietary strains, such as CNCM I-1079, CNCM I-3799, and DBVPG 6763. The genomic and functional equivalence of these strains is frequently assumed but remains insufficiently characterised. This strain-level heterogeneity can raise concerns about reproducibility, regulatory oversight, and whether strain-specific traits contribute to differences in clinical efficacy and the development of advanced microbiome therapeutics (14–16).

Despite their shared ancestry between *S. cerevisiae* and *S. boulardii*, direct comparative analyses have been limited in depth and often rely on a single *S. cerevisiae* strain, failing to account for strain-level diversity within the species. This gap has contributed to overstating the distinctiveness of *S. boulardii* and underestimating the probiotic potential inherent to certain *S. cerevisiae* strains.

In this study, we compared five commercially available *S. boulardii* strains from different manufacturers with three *S. cerevisiae* strains, including two commercial baker’s yeasts and one laboratory strain, CEN.PK113-5D. We analysed genomic variation, growth dynamics under stress-relevant conditions, and metabolic profiles to identify probiotic-relevant traits. We further evaluated potential health benefits of each strain and addressed misconceptions regarding the growth properties of *S. boulardii* relative to *S. cerevisiae*. Together, these findings provide a comparative framework for defining yeast-based probiotic functionality and guiding the selection or engineering of next-generation probiotic yeasts.

## Materials and Methods

### Strain isolation and culture condition

*Saccharomyces cerevisiae* strains and *Saccharomyces boulardii* strains were isolated from commercial products (Table 1) by streaking onto YPD agar plates and incubating at 37°C for 48 hours. Individual colonies were re-streaked twice to ensure clonal purity. For strains retrieved from cryostock, cultures were revived on YPD agar plates at 37°C for 72 hours and stored at 4°C for no longer than two weeks prior to use. Single colonies were precultured overnight in one of the following media prior to experimentation: defined minimal medium (DMM; pH 6) composed of 7.5 g/L (NH_4_)_2_SO_4_, 14.4 g/L KH_2_PO_4_, 0.5 g/L MgSO_4_·7H_2_O, 20 g/L glucose, 2 mL/L trace metals solution, and 1 mL/L vitamins(17), yeast extract peptone dextrose (YPD; pH 6) medium containing 10 g/L yeast extract, 20 g/L casein peptone, and 20 g/L glucose (Sigma-Aldrich), or de Man, Rogosa and Sharpe (MRS; pH 6.5) medium (Sigma-Aldrich Cat. No. 1.10661.0500).

**Table 1.** Origin, commercial brand, and identification of *S. cerevisiae* and *S. boulardii* strains used in this study.

| Strain origin | Brand | Strain | Strain ID |
| --- | --- | --- | --- |
| Biocodex | Precosa | *S. boulardii* | Sb CNCM I-745 |
| Gnosis by Lesaffre | Sachhaflor | *S. boulardii* | Sb DBVPG 6763 |
| ATCC | MYA-796 | *S. boulardii* | Sb ATCC MYA-796 |
| Gnosis by Lesaffre | NOW Foods | *S. boulardii* | Sb CNCM I-3799 |
| Lallemand | Invivo Bio.Me | *S. boulardii* | Sb CNCM I-1079 |
| KronJäst | KronJäst för Matbröd | *S. cerevisiae* | Sc baker's yeast Swe |
| Lallemand | Malteserkorsgær | *S. cerevisiae* | Sc baker's yeast DK |
| Euroscarf | N/A | *S. cerevisiae* | Sc CEN.PK113-5D |

### High molecular weight yeast DNA extraction

Yeast cells were initially cultured in 25 mL DMM in baffled shake flasks for 24 hours. Cells were subsequently harvested by centrifugation and resuspended to achieve an optical density 600 nm (OD_600_) of approximately 30, corresponding to approximately 3 × 10^8^ cells per aliquot, in DNA/RNA Shield (Zymo Research, Cat. No. R1100) and stored at 4°C prior to DNA extraction. DNA extraction was performed according to the protocol previously described(18).

### DNA sequencing, assembly and gene calling

Genomic DNA from eight Saccharomyces strains was sequenced by Plasmidsaurus using Oxford Nanopore Technologies and processed using a standardised assembly and annotation pipeline. Raw reads were quality filtered using Filtlong v0.2.1, discarding the lowest 5% of reads based on a composite Phred quality and length score (with --mean_q_weight 10). High-quality reads were assembled de novo with Flye v2.9.1, using parameters optimised for long read sequencing input. Contigs were annotated with Augustus v3.5.0, using an automatically selected gene model based on the closest reference genome. Open reading frames (ORFs) were queried against the UniProt v2024_04 database using BLAST v2.15.0, and top hits with e-values < 0.05 were retained. Assembly structure was visualised using Bandage v0.8.1, and genome completeness was assessed with BUSCO v5.8.0.

### Pan genome construction and comparative genomics

A pan-genome was constructed for all eight strains using Anvi’o v8 (19), incorporating external gene calls generated by Augustus. Contig databases were annotated with Pfam v37.4 domain families and KEGG orthologs, then merged into a unified genome storage.

Gene clusters were identified via an all-vs-all DIAMOND search, followed by clustering using the Markov Cluster Algorithm with an inflation parameter of 2.0 and a minimum bit-score ratio of 0.5. Clusters were classified as core (present in all genomes), accessory (present in multiple but not all genomes), or singleton (unique to one genome). Comparative analysis identified gene clusters uniquely shared among S. cerevisiae or S. boulardii strains. Pairwise average nucleotide identity (ANI) was computed using FastANI (20) to assess overall genomic similarity.

### K-mer composition

To evaluate compositional bias across strains, two k-mer data sets were compiled. (i) Whole-genome 3-mers: Counts were taken directly from each assembly. For every k-mer, its reverse complement was computed, and the pair was collapsed to a single canonical bin defined as the lexicographically smaller sequence; counts from both orientations were summed, yielding 32 canonical trinucleotides. (ii) Coding-sequence counts were obtained by extracting every CDS feature from the GenBank files with Biopython 1.81, writing each exon in its translated orientation, and concatenating the fragments per strain; complements were not merged so strand specificity was retained.

Three-mers (k=3) were enumerated with Jellyfish 2.3.0 (21), producing a frequency table of the 64 possible codons for each strain. To make profiles comparable across genomes of different length, every column was converted to relative frequency

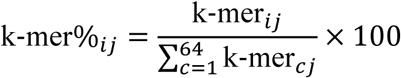

Where k-mer_ij_ is the raw count of k-mer *i* in strain *j*, and the denominator sums the counts of all 64 codons (c = 1 to 64) in strain *j*.

The resulting whole-genome and CDS percentage matrices served as input for downstream statistical comparisons and visualisation.

### Whole genome synteny and structural variation

Whole-genome synteny was assessed by aligning the eight yeast assemblies to the *Saccharomyces cerevisiae* S288C reference with Minimap2 v2.17, which produces base-level SAM alignments for structural analyses. The resulting SAM files were converted to coordinate-sorted BAM format with Samtools v1.10, for downstream processing. To enable chromosome-scale comparison between strains, we generated 16-chromosome pseudo-genomes for each query with Chroder (bundled with SyRI), which concatenates contigs according to their reference coordinates, and realigned these pseudo-chromosomes to the reference with the same Minimap2 settings. Syntenic blocks and structural rearrangements (inversions, translocations, duplications and indels) were identified with SyRI v1.5 (22) executed on the sorted BAMs. Finally, synteny maps for all strains were visualised with Plotsr v0.2which renders a vertical plot of reference chromosomes against each pseudo-genome, colouring collinear blocks and exposing structural breakpoints. All tools were run with default parameters.

### Growth assessments

*S. cerevisiae* and *S. boulardii* strains were inoculated at OD_600_ 0.1 in 200 µl media in CELLSTAR 96-well microtiter plates (Greiner Bio-One, Cat. No. 655160) with a Breathe-Easy sealing membrane (Sigma Aldrich, Cat. No. Z380059). OD_600_ was measured using a bench-top spectrophotometer. Growth was monitored at 30°C, 37°C, or 42*°*C in DMM, YPD, or MRS, respectively. For acid tolerance assay, DMM and YPD were adjusted to pH 1.5, 2.5, 3.5, or 4.5 by dropwise addition of HCl prior to inculcation. Anaerobic conditions were established by streaking and preculture strains in sealed containers with AnaeroGen pouches (Thermo Fisher Scientific, Cat. No. AN0035A) followed by incubation in microtiter plates in a Coy Anaerobic Chamber (gas mixture, 95% N_2_ and 5% H_2_). All media used for anaerobic growth was as pre-reduced for 48 hours. Real-time OD_600_ was measured every 15 min for 48 hours using a BioTek Synergy H1 microplate reader (Agilent Technologies).

### Growth rate estimation

OD_600_ time-series data were collected from microplate cultures and processed in R and converted to natural logarithms after filtering out non-positive values. For each condition (strain × medium × temperature × oxygen × replicate), we estimated the maximum specific growth rate (μ_max) using a rolling-window linear regression approach. Briefly, linear models of log(OD_600_) versus time (in hours) were fitted across all contiguous windows of 4–5 data points. For each window, the slope of the regression (h^-1^) and the corresponding coefficient of determination (R^2^) were recorded. The window yielding the highest slope (with R^2^ used as a tie-breaker) was selected as the best estimate of exponential growth. The corresponding μ_max (h^-1^) and doubling time (dt = ln(2)/μ_max, reported in both hours and minutes) were calculated.

### Viability assessments

Pre-cultures were diluted to OD_600_ of 10 in a final volume of 500 μL using artificial intestinal fluid (pH 6.8; Biochemazone, Cat. No. BZ176), artificial colonic fluid (pH 7.0; Biochemazone, Cat. No. BZ177), artificial gastric fluid (pH 1.5; Biochemazone, Cat. No. BZ175), or 0.1 M H_2_0_2_ prepared in YPD and incubated at 37°C with shaking. At 0, 3, 6, 24, and 48 hours, 10 µL of culture was harvested and diluted in 90 μL filtered PBS in a clear-bottom microplate, to achieve an approximate OD_600_ of 1. Cell viability was assessed using the Yeast Live-or-Dye™ Fixable Live/Dead Staining Kit (Biotium, Cat. No. 31064) according to the manufacturer’s instructions. Samples were analysed using a Novocyte Quanteon (Agilent Technologies). Forward scatter (FSC) and side scatter (SSC) were measured with a gain of 400; non-viable cells were detected through PE detection channel with a 586 nm filter (yellow laser) and a gain of 591. A threshold of 6000 was used, and data acquisition continued until 30 μL of the sample was injected. Gates for dead cells identification were defined using heat-killed controls (100°C for 10 min) treated under identical exposure conditions.

### HPLC analysis of yeast metabolites

Strains were cultured in triplicate by diluting overnight cultures to OD_600_ 0.1 in either 500 µL of DMM, YPD, or MRS in 96 deep-well plates, or 25 mL of DMM in 250 mL baffled shake flask. Aerobic cultures were incubated at 37°C under 250 rpm shaking. Anaerobic cultures were incubated in a sealed container with AnaeroGen pouches at 37°C. Yeast supernatants were obtained by centrifuging cultures at 4,000 × g for 10 minutes at 8, 24, 48, 72, and 120 hours. The supernatants were then stored at –20 °C until further analysis.

(D+)-Glucose monohydrate (Sigma Aldrich, Cat. No. 16301), ethanol absolute (VWR, Cat. No. 20821.310), sodium acetate (Sigma Aldrich, Cat. No. S8750), sodium L-lactate (Sigma Aldrich, Cat. No. 71718), and sodium succinate dibasic (Sigma Aldrich, Cat. No. 14160) were quantified by high-performance liquid chromatography (HPLC) using a Dionex Ultimate 3000 HPLC system with analysis software Chromeleon (Thermo Fisher Scientific). Metabolites were separated on Bio-Rad Aminex HPx87 column with 5 mM H_2_SO_4_ as an eluent at a flow rate of 0.6 mL/min. Column temperature was maintained at 30°C, and detection was performed using a refractive index detector. The limit of detection for all metabolites was 0.1 g/L.

### Mammalian cell culture maintenance

The human intestinal epithelial cell line HT-29 (ATCC Cat. No. HTB-38) and the HT29-Lucia AhR (aryl hydrocarbon receptor) reporter cell line (InvivoGen, Cat. No. ht2l-ahr) were cultured in McCoy’s 5A (Modified) medium (Gibco, Thermo Fisher Scientific, Cat. No. 16600082) supplemented with 10% foetal bovine serum (FBS; Thermo Fisher Scientific, Cat. No. A5670701) and 1% penicillin–streptomycin (Thermo Fisher Scientific, Cat. No. 15140148). Cells were maintained in a humidified incubator at 37°C with 5% CO₂.

Human THP-1 monocytes (ATCC, Cat. No. TIB-202) were cultured in RPMI-1640 medium (Thermo Fisher Scientific, Cat. No. 12027599) 10% FBS and 1% penicillin–streptomycin and incubated at 37°C with 5% CO₂. For macrophage differentiation, cells were seeded at appropriate density and treated with 20 µM phorbol 12-myristate 13-acetate (PMA; Sigma Aldrich, Cat. No. P8139) for 24 hours, followed by washing with serum-free RPMI-1640 prior stimulation.

HT29-Lucia AhR reporter cells were thawed and passaged twice in the absence of the selection antibiotic, then maintained in compete McCoy’s 5A medium supplemented with 100 µg/mL Zeocin (Invivogen; Cat. No. 11006-33-0) and 1% penicillin–streptomycin. Cells were harvested prior to reaching 90% confluency.

THP-1-Lucia NF-κB (Nuclear factor kappa B) cells (ATCC Cat. No. TIB-202-NFkB-LUC2) were maintained in suspension and were not differentiated with PMA prior to use. Cells were cultured in RPMI-1640 media containing 1 µg/ml puromycin (Thermo Fisher Scientific, Cat. No. #A1113803), 10% FBS, and 0.05 mM 2-Mercaptoethanol (Thermo Fisher Scientific, Cat. No. #21985023).

All cell lines were passaged every 2–3 days and used for experiments within 10 passages. All experiments were performed using three to five biological replicates.

### Preparation of yeast supernatant

For the preparation of yeast supernatants, strains were diluted to an initial OD_600_ of 0.1 in 25 mL of DMM in 250 mL shake flasks and incubated at 37°C with shaking at 250 rpm for 24 hours. Cultures were prepared in duplicate and pooled at the end of the incubation period. The final OD_600_ was measured using a bench-top spectrophotometer to ensure similar biomass formulation across strains. Cultures were then centrifuged at 4,000 × g for 10 minutes, and the resulting supernatants were sterile-filtered through 0.22 μm syringe filters. Filtered supernatants were aliquoted and stored at –20 °C and were thawed only once prior to use.

### LPS and TNFα stimulation of mammalian cells

For HT-29 stimulation, 10,000 cells were seeded in 100 μL complete McCoy’s 5A medium in CELLSTAR 96-well microtiter plate and incubated for 48 hours to reach 90% confluency. Subsequently, the media were replaced with 200 μL of fresh McCoy’s 5A media containing 1 μg/mL lipopolysaccharides (LPS from *Escherichia coli* O111:B4; Sigma Aldrich, Cat. No. L2630) or 10 ng/mL tumour necrosis factor-α (TNFα; R&D Systems, Cat. No. 10291-TA), supplemented with 10% (v/v) yeast supernatant. Control groups received 10% (v/v) of DMM. Supernatants were collected at 6- or 24-hours post-stimulation.

For THP-1 stimulation, 50,000 cells were seeded in 100 μL complete RPMI*-*1640 medium with 5 µg/ml PMA in CELLSTAR 96-well microtiter plate and incubated until a 90% confluent monolayer formed. Subsequently, the cells were replaced with 200 μL of fresh RPMI-1640 media containing 1 μg/mL LPS or 10 ng/mL TNFα and 10% (v/v) yeast supernatant. Control groups received 10% (v/v) of DMM. Supernatants were collected at 24-hours post-stimulation.

### Cytokine quantification by ELISA

Cytokine concentrations in the supernatants from stimulated HT-29 and THP-1 cells were quantified by ELISA. Interleukin-8 (IL-8) was measured using the DuoSet ELISA kit (R&D systems, Cat. No. DY208), according to the manufacturer’s protocol. TNFα was quantified using the TNFα ELISA kit (Abcam, Cat. No. ab181421). Absorption at 450nm was measured using a BioTek Synergy H1 microplate reader (Agilent Technologies).

### NF-**κ**B activation reporter assay in THP1 NF-kB-Luc2 cells

THP1 NF-kB-Luc2 reporter cells were seeded at 100,000 cells per well in 200 μL complete RPMI*-*1640 medium in CELLSTAR 96-well microtiter plates. Cells were stimulated with 100 ng/mL LPS or 10 ng/mL TNFα and 10% (v/v) yeast supernatant. Control groups received 10% (v/v) of DMM. After 6 hours incubation at 37°C with 5% CO2, 100 µL of samples from each of the wells were transferred to a white 96-well microtiter plate (Thermo Fisher Scientific, Cat. No. 165305) and 100 µL of the Bright-Glo Luciferase Assay reagent (Promega, Cat. No. E2610), prepared according to the manufacturer’s protocol, was added to the wells. Luminescence was immediately measured on a BioTek Synergy H1 microplate reader (Agilent Technologies).

### AhR activation reporter assay in HT29-Lucia cell

HT29-Lucia AhR reporter cells were seeded at 50,000 cells per well in 180 µL medium in CELLSTAR 96-well microtiter plates. After adherence, 20 µL of yeast supernatant, control DMM, or 20 µM 6-formylindolo[3,2-b]carbazole (FICZ; Sigma-Aldrich, Cat. No. 172922-91-7) was added per well. Cells were incubated for 48 hours at 37°C. Following incubation, 20 μL of cell culture supernatant was transferred to a black 96-well plate with optical bottom. Luminescence was detected by adding 50 μL QUANTI-Luc (Invivogen, Cat. No. rep-qlc4r2) and immediately measured using a BioTek Synergy H1 plate reader.

### TEER analysis of differentiated Caco-2 monolayers co-cultured with live *S. boulardii*

Caco-2 cells were seeded at 20,000 cells per well on 24-well Transwell inserts (0.4 µm pore size; 0.33 cm^2^ PET membrane; cellQART, Cat. No. #9320402) and differentiated for 19–21 days in DMEM (Thermo Fisher Scientific, Cat. No. #11965092) supplemented with 10% FBS and 1% penicillin–streptomycin (Thermo Fisher Scientific, Cat. No. #15140122). Media were replaced every other day during differentiation. On the day of the assay, the apical compartment was refreshed with 150 µL of live *S. boulardii* suspension (1×10^8^ cells/mL; OD_600_ = 1) in DMEM with 10% FBS and 1% penicillin–streptomycin; the basolateral chamber received 600 µL of cell free matching medium. Transepithelial electrical resistance (TEER) was recorded at multiple time points over 24 hours using a Millicell ERS-2 voltohmmeter (Merck, Cat. No. MERS00002) with an STX-04 electrode. Percentage change in TEER at each time point was calculated relative to the 0h post-treatment baseline. The area under the curve (AUC) was determined over the 24-hour period to quantify the barrier integrity changes.

### Data analysis

All experiments were performed with three to five biological replicates and compiled from one to two independent experimental runs. All statistical analysis was performed using Graph Pad Prism software (Graph Pad Software Inc.) or RStudio version 4.1.0, utilising the rstatix and DescTools packages. Data are presented as mean ± standard error of the mean (SEM). Statistical analyses were conducted using one-way ANOVA followed by either Dunnett’s or Tukey’s post hoc tests, as appropriate, to account for multiple comparisons. In addition, independent two-sample t-tests were performed with false discovery rate (FDR) correction for multiple testing where applicable. A *P* value of < 0.05 was considered statistically significant.

## Results

### Comparative genomic analysis reveals conserved and divergent features distinguishing *S. boulardii* from *S. cerevisiae*

To assess the genetic difference between *S. boulardii* and *S. cerevisiae*, we performed whole-genome comparisons of the five representative *S. boulardii* isolates and three *S. cerevisiae* strains. All *S. boulardii* isolates exhibited high sequence conservation, with >99.98% average nucleotide identity (ANI), a genome size of 11.8 Mb, and ∼6,400 predicted genes (Fig. 1A; Supplementary Table S1). While the overall genomic similarity between *S. cerevisiae and S. boulardii* was also high (ANI >99%), *S. cerevisiae* genomes were larger (∼12.3 Mb) and each contained over 100 singletons, indicating broader functional diversity of *S. cerevisiae* strains. We later delineated gene clusters unique to either *S. boulardii* or *S. cerevisiae* (“*S. boulardii* only” and “*S. cerevisiae* only”; Fig. 1A; Supplementary Table S2). Both strain-specific sets were sparsely annotated, reflecting limited coverage of these open reading frame in existing functional databases. Alternatively, these clusters may encode recently evolved or highly diverged genes lacking homologs in current reference collections.

**Figure 1.**
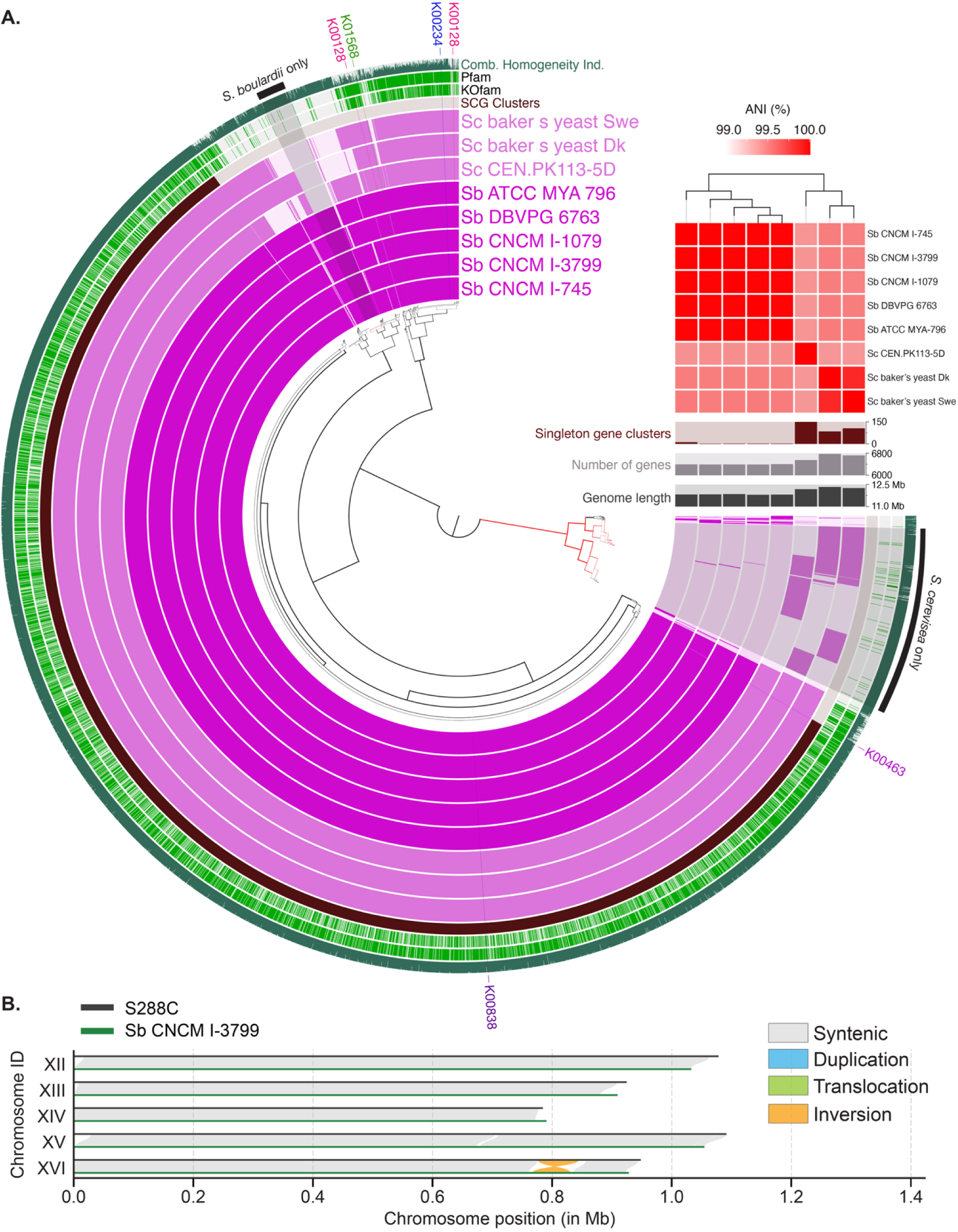
Comparative genomic analysis of *Saccharomyces* strains. **(A)** Pangenome analysis of eight Saccharomyces strains using anvi’o. The central dendrogram reflects hierarchical clustering of gene clusters based on their presence or absence across genomes. Each concentric layer corresponds to a single genome, with coloured bars marking the presence of a given gene cluster. Clusters are classified as single-copy core genes (SCGs; present in all genomes), accessory (present in 2–7 genomes), or singletons (strain-specific). Green rings denote clusters with functional annotations in either KOfam or Pfam databases. The outermost ring shows the combined homogeneity index, which summarizes sequence conservation across genomes within each cluster based on amino acid identity. **(B)** Whole-genome alignment of *S. boulardii* CNCM I-3799 to the *S. cerevisiae* reference strain S288C reveals a conserved 68.3 kb inversion on chromosome XVI (highlighted in orange). Grey blocks indicate syntenic regions. This inversion was present in all *S. boulardii* strains and absent in all *S. cerevisiae* strains (see Supplementary Figure S2 for other isolates).

*S. boulardii* harboured a distinct set of conserved genes across *S. boulardii* isolates (Supplementary Table S2). Only three of these clusters contained Pfam-annotated domains: ribosomal protein S19, a Flo11 adhesin domain, and a cytochrome c/quinol oxidase polypeptide. The presence of a Flo11 adhesin domain could suggest a potential role in adhesion or host interaction, although the gene encodes only a partial domain and lacks full-length homology to canonical *FLO11*.

While gene content was broadly similar between *S. boulardii* and *S. cerevisiae*, differences in gene copy number may also contribute to strain-specific phenotypes. Analysis of carbohydrate-active enzyme (CAZyme) copy numbers revealed *S. cerevisiae* generally harboured more CAZyme-encoding genes than *S. boulardii*, except for AA4 and AA7 families, encoding vanillyl-alcohol and glucooligosaccharide oxidases, which were enriched in *S. boulardii* (Supplementary Table S3). These enzymes may play roles in the oxidative metabolism of aromatic compounds and oligosaccharides and could contribute to lineage-specific metabolic capabilities. We also observed increased representation of enzymes from 12 distinct classes in *S. boulardii* compared to *S. cerevisiae*, including glucan 1,3-β-glucosidases and acid phosphatases (Supplementary Table S4). These enzymes may contribute to differences in membrane architecture (23) and probiotic functionality among the strains (24).

Chromosomal alignment also revealed conserved synteny between *S. boulardii*, with no large-scale translocations. However, a 68,263 bp inversion (position 776,149–844,412) on chromosome XVI was consistently observed in all *S. boulardii* isolates (Fig. 1B; Supplementary Fig. S1), distinguishing them from *S. cerevisiae*. To obtain alignment-free view of sequence divergence, we also quantified genome-wide k-mer frequencies. This “genomic signature” collapses SNPs, indels and horizontally transferred fragments into a single high-dimensional profile. In a study of >260 *S. cerevisiae* isolates, this method resolves population structure as accurately as SNP-based methods while capturing additional diversity stemming from insertions and horizontal transfers, highlighting the method’s strain level resolution (25). Genome wide patterns of sequence composition across all strains were largely conserved (Supplementary Fig. S2A), with modest strain-level variation but no consistent differences separating *S. boulardii* from *S. cerevisiae*, reflecting their close phylogenetic relationship. We next examined codon-usage frequencies across strains using annotated coding sequences (Supplementary Fig. S2B). While clustering did not perfectly resolve the two species, *S. boulardii* strains tended to group separately from *S. cerevisiae*, suggesting subtle but lineage-associated codon preferences. Strain-level variation was evident within each species, yet certain codons exhibited consistent differences—for example, the ATT codon encoding isoleucine was used less frequently in *S. boulardii* than in *S. cerevisiae* strains.

To identify metabolic features distinguishing *S. boulardii* from *S. cerevisiae*, we compared protein-coding variants in central metabolism and tryptophan catabolism across the five *S. boulardii* and three *S. cerevisiae* genomes. All *S. boulardii* isolates retained the *SDH1^H202Y,F318Y^* double substitution in succinate dehydrogenase, previously linked to elevated acetate production (26), confirming its conservation across the lineage (Supplementary Table S5). Given *S. boulardii*’s ability to activate the aryl hydrocarbon receptor (27), we next surveyed the two upstream tryptophan-catabolic enzymes: indoleamine 2,3-dioxygenase (*BNA2*) and tryptophan aminotransferase (*ARO8*). *BNA2* harboured two *S. boulardii*-specific substitutions (T268S and R371K) absent in all *S. cerevisiae* strains, and *ARO8* contained a conserved E495G substitution (Supplementary Table S6–7). In contrast, downstream enzymes, pyruvate decarboxylase (*PDC1*) and the aldehyde dehydrogenases (*ALD6*/*ALD5*), showed no unique variants in *S. boulardii* (Supplementary Table S8–10). These lineage-specific changes in *BNA2* and *ARO8* may redirect flux toward indole derivatives, providing a mechanistic basis for AhR activation by *S. boulardii*.

### *S. boulardii* exhibit enhanced growth and viability under host-relevant conditions

Building on our genomic insights, we next evaluated *S. boulardii*’s phenotypic performance under host-relevant conditions. We measured proliferation and survival in defined minimal and rich media across a range of temperatures, oxygen levels, pH values, bile concentrations and simulated gastrointestinal fluids to model both *in vitro* and *in vivo* stressors.

To assess thermotolerance at host-relevant temperatures, we compared the growth of *S. boulardii and S. cerevisiae* in defined minimal medium (DMM) and rich media (YPD, MRS) at 30°C, 37°C and 42°C under aerobic conditions. At both 30°C and 37°C, maximum specific growth rates were comparable between species; however, under the different conditions tested, most *S. cerevisiae* strains grew 20–80% faster than *S. boulardii* (with CEN.PK113-5D matching *S. boulardii* kinetics) (Fig. 2A; Supplementary Fig. S3A–L). Neither species proliferated at 42°C, indicating that both yeasts lack thermotolerance above this temperature (Supplementary Fig. S3C).

**Figure 2.**
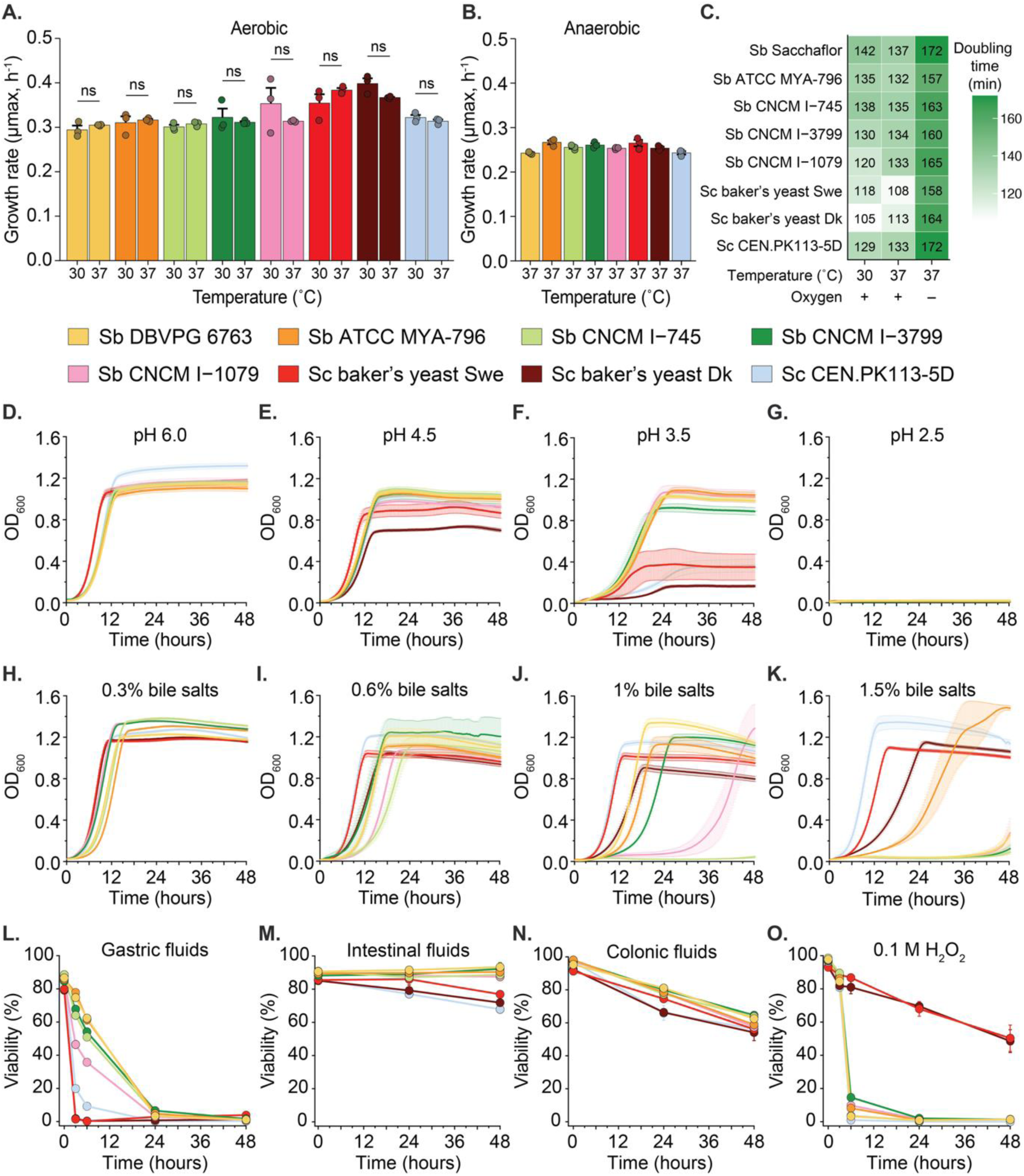
Growth and survival characterisation of *S. cerevisiae* and *S. boulardii* strains under host-relevant conditions. Growth rates measured over 48 hours at **(A)** 30°C and 37°C under aerobic conditions, and **(B)** at 37°C under anaerobic conditions. **(C)** Mean doubling times (in minutes) calculated for cultures grown aerobically at 30°C and 37°C, as well as anaerobically at 37°C. Growth curves in DMM adjusted to pH **(D)** 6.0, **(E)** 4.5, **(F)** 3.5, and **(G)** 2.5 under aerobic conditions at 37°C. Growth curves in DMM supplemented with **(H)** 0.3%, **(I)** 0.6%, **(J)** 1.0%, and **(K)** 1.5% bile salts, under aerobic conditions at 37°C. Viability following 48-hour exposure to **(L)** artificial gastric fluid, **(M)** artificial intestinal fluid, **(N)** artificial colonic fluid, and **(O)** 0.1 M hydrogen peroxide (H_2_O_2_) in PBS was measured by flow cytometry. All data represent mean + or ± SEM from three independent biological replicates. Statistical significance was determined using independent two-sample t-tests, with multiple comparisons adjusted using FDR correction. Significance was accepted at *P* < 0.05 (\**P* < 0.05, \*\**P* < 0.01).

To mimic the anaerobic gut environment, we measured growth at 37°C under oxygen limitation. All strains grew anaerobically; however, *S. cerevisiae* outpaced *S. boulardii* in rich media (Supplementary Fig. S4), whereas both species grew at comparable rates in DMM (Fig. 2B). In DMM, anaerobic growth rates fell 20–45% below aerobic rates (P < 0.05, two-sample t-test with FDR correction) (Fig. 2C; Supplementary Fig. S4E), and *S. boulardii* experienced a larger reduction in proliferation than *S. cerevisiae* in rich media under anaerobiosis.

Next, to evaluate tolerance to gastrointestinal stresses, we cultured strains in DMM and YPD at pH 1.5–4.5 or with 0.3–2.5% bile salts. *S. boulardii* sustained growth down to pH 3.5 in DMM and pH 2.5 in YPD, whereas *S. cerevisiae* growth declined sharply below pH 3.5 in DMM and 4.5 in YPD (Fig. 2D–G; Supplementary Fig. S5A–D). Conversely, *S. cerevisiae*, including CEN.PK113-5D, which showed no defect at 2.5% bile, outperformed *S. boulardii* in bile resistance (Fig. 2H–K; Supplementary Fig. S5E). Thus, *S. boulardii* is more acid-tolerant but less bile-resistant than *S. cerevisiae*, reflecting potential distinct survival strategies in the upper gastrointestinal tract.

Finally, we quantified cell viability after exposure to simulated gastric, intestinal and colonic fluids, as well as to 0.1 M hydrogen peroxide (H_2_O_2_)–induced oxidative stress, using Live/Dead staining and flow cytometry (Supplementary Fig. S5F–G). *S. boulardii* exhibited markedly higher survival in both gastric and intestinal fluids compared to *S. cerevisiae* strains (Fig.s 2L– M), with approximately 50% viability retained after 6 hours in gastric fluids, whereas *S. cerevisiae* strains showed 0–10% survival. In contrast, no major differences in viability were observed between strains upon exposure to colonic fluids (Fig. 2N). Under oxidative stress, commercial baker’s yeasts exhibited higher survival than both *S. boulardii* and CEN.PK113-5D, suggesting that enhanced catalase or peroxidase activity confers superior resistance to H_2_O_2_ (Fig. 2O).

### *S. cerevisiae* and *S. boulardii* exhibit distinct central carbon metabolism profiles across culture conditions

Having characterised growth and stress responses, we next examined how these strains channel carbon through fermentation and respiration. We profiled biomass accumulation, substrate uptake and byproduct formation over a 72-hour time course in 250 mL shake-flask cultures.

In shake flasks, optical density 600 nm (OD_600_) increased more rapidly in commercial baker’s yeast strains during the first 24 hours, but each strain reached its own plateau by 72 hours (Fig. 3A). Glucose depletion proceeded fastest in baker’s yeasts, 50% of initial glucose was consumed within 8 hours, whereas *S. boulardii* strains utilised glucose more slowly and completed depletion by 24 hours (Fig. 3B). Ethanol accumulation peaked at 24 hours for all strains; baker’s yeasts attained higher maximum ethanol titters than *S. boulardii*, and all strains then re-consumed ethanol during the subsequent respiratory phase (Fig. 3C).

**Figure 3.**
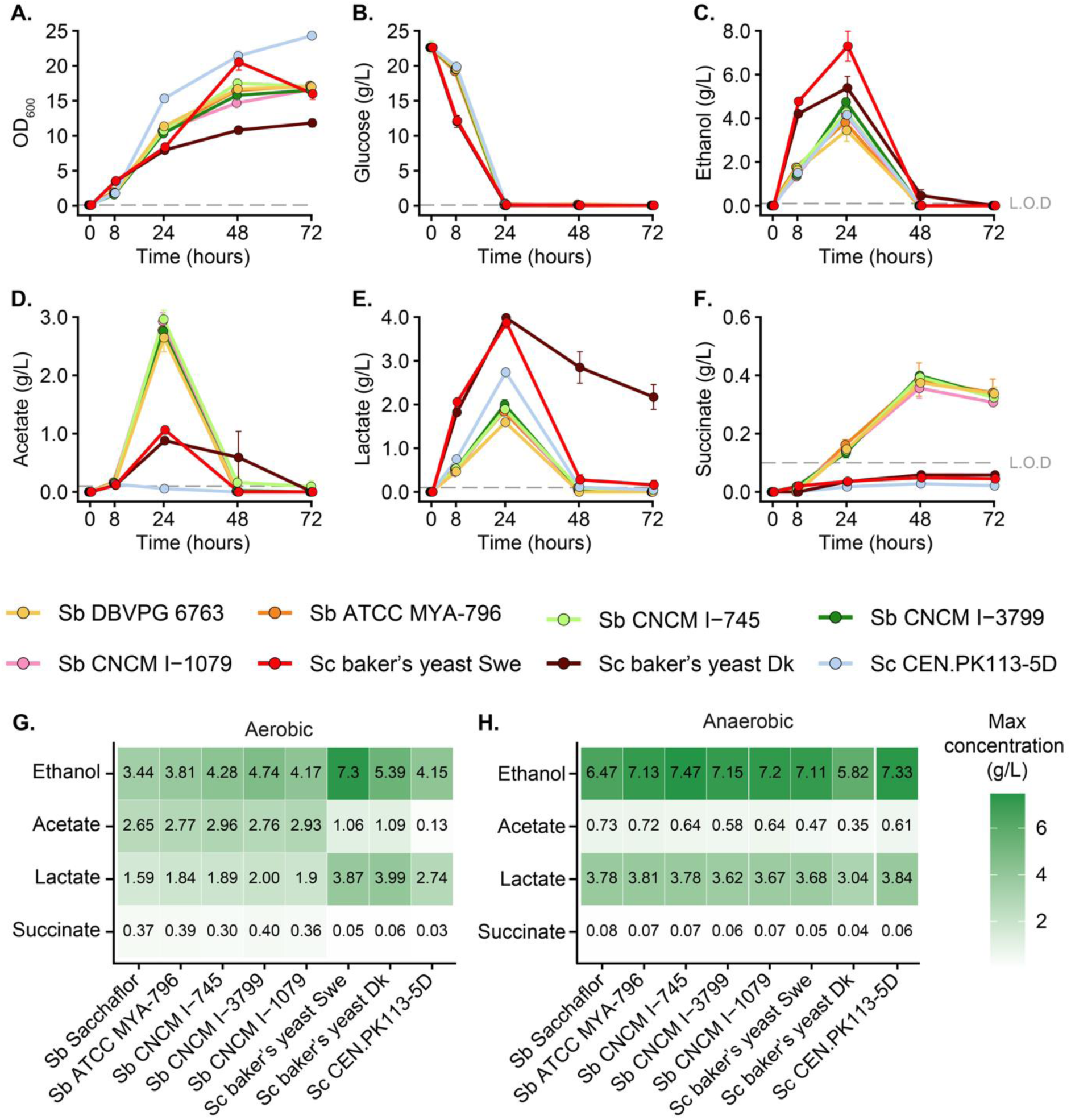
Time-course profiling of metabolite production by *S. cerevisiae* and *S. boulardii* strains. **(A)** Biomass accumulation, measured as OD_600_, was monitored at 0, 8, 24, 48, and 72 hours. Concentrations of key extracellular metabolites, **(B)** glucose, **(C)** ethanol, **(D)** acetate, **(E)** lactate, and **(F)** succinate, in the culture supernatants were quantified at 0, 8, 24, 48, and 72 hours. Maximum metabolite concentrations under **(G)** aerobic cultivation conditions across 72-hour time points and **(H)** anaerobic cultivation conditions across 120-hour time points. Data represent mean values from three independent biological replicates ± SEM.

To assess redox balancing via organic acids, we measured acetate, lactate, and succinate. *S. boulardii* produced ∼3 g/L acetate by 24 hours, compared to ∼1 g/L in baker’s yeasts; CEN.PK113-5D generated only trace acetate (Fig. 3D). Lactate accumulation reached ∼4 g/L in baker’s yeasts versus ∼2 g/L in *S. boulardii* (Fig. 3E). Both acetate and lactate declined after 48 hours, indicating their further oxidation. In contrast, succinate levels were consistently higher in *S. boulardii* (Fig. 3F) and remained stable through 72 hours, suggesting its role as a terminal respiratory byproduct.

We then explored whether culture format alters these metabolic patterns by repeating the experiment in 96-deep-well plates under aerobic conditions in DMM, YPD, and MRS. Although glucose depletion dynamics resembled those in shake flasks, ethanol persisted up to 120 hours and acetate and lactate remained detectable, reflecting slower respiratory turnover due to limited aeration (Fig. 3G–H; Supplementary Fig. S6).

Finally, complete anaerobiosis further highlighted oxygen’s role: glucose consumption extended to 120 hours, with baker’s yeast strains still showing modestly faster uptake (Supplementary Fig. S7). Ethanol, acetate and lactate levels plateaued in both species, consistent with constrained oxidative metabolism and minimal post-fermentative turnover. As a result, the metabolic distinctions observed under aerobic conditions disappeared under oxygen limitation.

### *S. boulardii* supernatant attenuate inflammatory responses

Building on our metabolic and growth characterisations, we next evaluated whether supernatant from *S. boulardii* modulates inflammation. To minimise confounding effects from undefined media components, all functional assays were performed using culture supernatants derived from strains grown exclusively in DMM. Supernatants were collected after 24 hours of shake-flask incubation—corresponding to the peak of acetate production— and subsequently applied to human epithelial and macrophage cell models to assess immunomodulatory effects.

To assess anti-inflammatory activity, we first used the human intestinal epithelial cell line HT-29, a well-established model for cytokine responses to TNFα stimulation. Supernatants from *S. boulardii* strains reduced TNFα-induced IL-8 secretion by ∼25% compared to TNFα alone (Fig. 4A), whereas *S. cerevisiae* supernatants had no effect. Neither *S. boulardii* nor *S. cerevisiae* supernatants significantly altered IL-8 or TNFα levels in response to lipopolysaccharides (LPS) stimulation (Fig. 4B–C), indicating stimulus-specific modulation.

**Figure 4.**
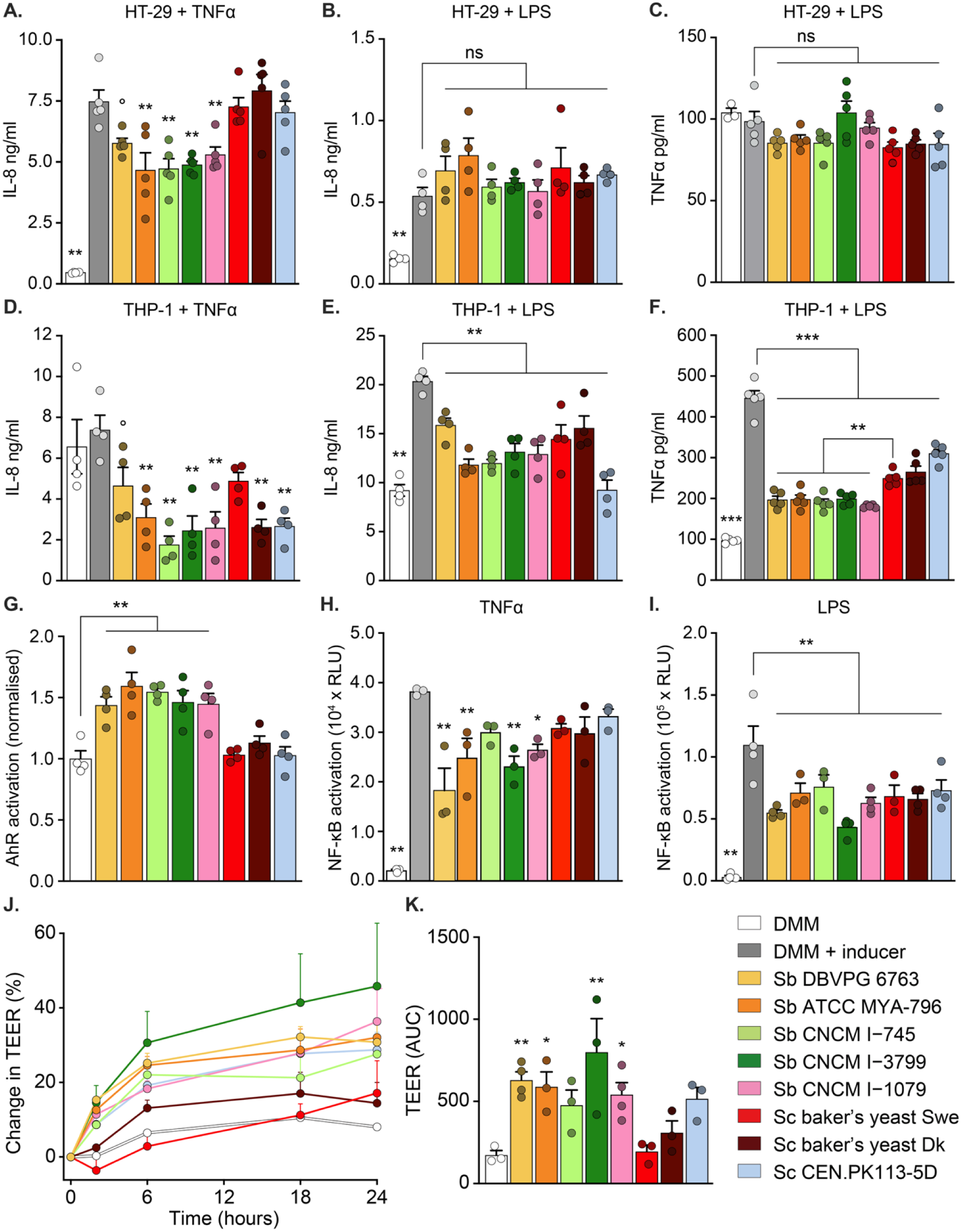
Immunomodulatory characterisation of *S. cerevisiae* and *S. boulardii* strains using human cell-based assays. **(A)** IL-8 production in HT-29 epithelial cells stimulated with TNFα (10 ng/mL) and treated with 10% (v/v) yeast supernatant for 24 hours. **(B)** IL-8 production in HT-29 cells stimulated with LPS (1 µg/mL) and treated with 10% (v/v) yeast supernatant for 24 hours. **(C)** TNFα production in HT-29 cells stimulated with LPS (1 µg/mL) and treated with 10% (v/v) yeast supernatant for 24 hours. **(D)** IL-8 production in THP-1 monocytic cells stimulated with TNFα (10 ng/mL) and treated with 10% (v/v) yeast supernatant for 24 hours. **(E)** IL-8 production in THP-1 monocytic cells stimulated with LPS (1 µg/mL) and treated with 10% (v/v) yeast supernatant for 24 hours. **(F)** TNFα production in THP-1 monocytic cells stimulated with LPS (1 µg/mL) and treated with 10% (v/v) yeast supernatant for 24 hours. **(G)** Normalised luciferase activity in AhR reporter cells following 24-hour exposure to 10% (v/v) yeast supernatant. **(H)** Luciferase activity in NF-κB reporter cells stimulated with TNFα (100 ng/mL) in the presence of 10% (v/v) yeast supernatant for 6 hours. **(I)** Luciferase activity in NF-κB reporter cells stimulated with LPS (100 ng/mL) and exposed to 10% (v/v) yeast supernatant for 6 hours. **(J)** Transepithelial electrical resistance (TEER) changes in epithelial monolayers after exposure to live yeast strains over 24 hours. **(K)** Area under the curve (AUC) of TEER change over the 24-hour exposure period. Data represent mean + SEM from three to five independent biological replicates. White bars indicate unstimulated controls treated with 10% DMM, while grey bars indicate TNFα or LPS stimulated controls treated with 10% DMM. Statistical significance for all panels, except (F), was determined using one-way ANOVA followed by Dunnett’s post hoc test, with either stimulated or unstimulated DMM-treated controls as the reference. For panel (F), one-way ANOVA followed by Tukey’s Honestly Significant Difference (HSD) test was used to correct for multiple comparisons. Significance was considered at *P* < 0.05 (° P < 0.1, * *P* < 0.05, ** *P* < 0.01, *** *P* < 0.001).

We further validated these findings in THP-1 monocyte-derived macrophages stimulated with TNFα or LPS. TNFα stimulation did not elevate IL-8 levels relative to media controls; however, supernatants from several *S. boulardii* and *S. cerevisiae* strains reduced IL-8 below baseline (Fig. 4D). LPS stimulation significantly increased IL-8 and TNFα production, and all yeast supernatants suppressed these responses. *S. boulardii* supernatants produced a significantly greater reduction in LPS-induced TNFα than *S. cerevisiae* strains (Fig. 4E–F).

To investigate underlying mechanisms, we assessed AhR and NF-κB pathway activity using reporter cell lines. In HT29-Lucia AhR reporter cells, supernatants from all *S. boulardii* strains increased AhR-driven luciferase activity by approximate 50% relative to control, indicating the presence of AhR ligands (Fig. 4G). *S. cerevisiae* supernatants did not significantly activate AhR signalling. NF-κB activity was measured in THP1–NFκB–Luc2 cells following TNFα or LPS stimulation. Under TNFα stimulation, most *S. boulardii* supernatants significantly suppressed NF-κB–dependent luciferase activity, whereas *S. cerevisiae* strains had no significant effect (Fig. 4H). Following LPS stimulation, all yeast supernatants reduced NF-κB activity to a similar extent (Fig. 4I).

We next tested whether *S. boulardii* directly enhances epithelial barrier function. Differentiated Caco-2 monolayers were incubated with live yeast, and transepithelial electrical resistance (TEER) was used to assess barrier integrity. Most *S. boulardii* strains significantly increased TEER relative to untreated controls, indicating a tightening of the epithelial (Fig. 4J–K). In contrast, *S. cerevisiae* baker’s strains had no significant effect (Fig. 4K).

## Discussion

This study addressed two questions: (i) which functional traits distinguish *S. boulardii* from its close relative *S. cerevisiae*, particularly in the context of probiotic activity, and (ii) whether *S. boulardii* strains from diverse sources exhibit consistent phenotypes across physiologically relevant and stress-inducing conditions.

Despite *S. boulardii* and *S. cerevisiae* sharing high overall sequence identity (>99% ANI), comparative genomics revealed distinct features that may underlie functional divergence. *S. cerevisiae* genomes were larger and encoded a broader set of singleton gene clusters, whereas *S. boulardii* harboured a smaller genome with a consistent set of lineage-specific genes, many of which remain functionally uncharacterised. These unique loci may contribute to strain-specific traits relevant to probiotic activity. Similar genomic signatures distinguishing *S. boulardii* from laboratory and industrial *S. cerevisiae* strains have been previously reported (9, 28, 29) and further substantiated by phenotypic and comparative analyses showing adaptations of *S. boulardii* to host-associated environments (30). Together, these findings suggest that the lineage-specific gene clusters consistently found across *S. boulardii* isolates may underlie its probiotic functions (28) and should be experimentally characterised to understand their function. Although codon usage profiles were broadly similar across strains, subtle shifts in synonymous codon preferences suggest adaptive responses to industrial propagation (31, 32), with potential implications for gene expression and performance *in vivo*. Such variation should also be considered when engineering *S. boulardii* or *S. cerevisiae* as alternative microbial therapeutics (14–16), as codon bias can influence heterologous gene expression (33).

Systematic growth profiling across diverse environmental conditions revealed key physiological distinctions among strains. Consistent with prior studies (8, 34, 35), *S. boulardii* strains exhibited similar growth kinetics at 30°C and 37°C. These findings challenge earlier claims that *S. boulardii* grows optimally at 37°C (7, 10, 16), which appear to reflect misinterpretations of earlier data(8). The commercial *S. cerevisiae* baker’s strain grew more rapidly than *S. boulardii* at both temperatures. Notably, earlier reports of superior *S. boulardii* growth at 37°C often involved comparisons with auxotrophic *S. cerevisiae* strains lacking uracil, histidine, and tryptophan biosynthesis, in contrast to wild-type *S. boulardii* (8). Such auxotrophies have been shown to impair *S. boulardii* growth under non-selective conditions (36), likely confounding earlier assessments of relative growth performance. These findings highlight the impact of strain background and experimental context in evaluating yeast physiology.

Further characterisation confirmed that *S. boulardii* displays greater resistance to low pH, while *S. cerevisiae* strains tolerate higher bile salt concentrations as shown previously (8, 9, 34, 35). Although thermal tolerance was not directly assessed, none of the strains grew at 42°C. Enhanced stress resistance *in S. boulardii* has been linked to its robust, mannan-rich cell wall (23, 37), which may support its survival in the gastrointestinal environment and contribute to immune modulation. Notably, engineering strategies that increase mannan biosynthesis in both species have yielded strains with improved stress resilience (38), highlighting cell wall composition as a tuneable feature for probiotic optimization.

The metabolic profiles of *S. cerevisiae* and *S. boulardii* were among the most distinct features between strains. Consistent with previous reports, *S. boulardii* produced higher levels of acetic acid than *S. cerevisiae*, a phenotype associated with polymorphisms in *SDH1* (26), both of which were conserved across all screened *S. boulardii* strains. We also observed elevated succinate production by *S. boulardii*, which accumulated over time and may likewise reflect variation in *SDH1* (39, 40). Notably, growth and metabolic output were strongly influenced by oxygen availability. Although, the anaerobic physiology of *S. boulardii* remains poorly characterised, this provides one of the few direct comparisons of growth under strictly anaerobic conditions. Under anaerobiosis, both S. cerevisiae and S. boulardii exhibited similar growth kinetics in DMM, though at substantially reduced rates compared to aerobic conditions. Supporting earlier findings (26), acetate production by *S. boulardii* was minimal or undetectable under anaerobic conditions, in contrast to its aerobic profile.

These results have implications for understanding *S. boulardii*’s activity *in vivo*. Although the gut lumen is predominantly anaerobic, steep oxygen gradients exist near the epithelial surface (41–43), potentially allowing *S. boulardii* transient access to oxygen and localised acetate production. Whether this is sufficient to mediate therapeutic effects remains unclear. Interestingly, *S. boulardii* has also been shown to increase acetate production in microbial communities under anaerobic conditions (27), suggesting potential microbiota-mediated mechanisms of action.

The immunomodulatory activity of yeast was strongly context-dependent, varying with inducer, cell type, and inflammatory readout. Nevertheless, across all tested conditions, *S. boulardii* consistently exhibited equal or greater anti-inflammatory effects than *S. cerevisiae*, underscoring its enhanced therapeutic potential.

Consistent with prior observations (44), exposure of HT-29 epithelial cells to *S. boulardii* supernatant significantly reduced IL-8 secretion, even at low concentrations (10% v/v), indicating potent anti-inflammatory activity. While acetate may contribute to this effect; given its ability to signal through GPR43, inhibit NF-κB activation, and suppress histone deacetylase activity (45), the observation that *S. cerevisiae* supernatants also reduced IL-8 under some conditions suggests the involvement of additional, strain-specific secreted factors (23, 28, 46, 47). The elevated acetate production by *S. boulardii* under aerobic conditions may account for its stronger immunosuppressive effects relative to *S. cerevisiae*. *In vivo*, such effects could be amplified by cross-feeding interactions, where *S. boulardii*-derived acetate is metabolised by commensal bacteria into butyrate or other anti-inflammatory short-chain fatty acids (48).

A distinct observation was the selective activation of AhR signalling by *S. boulardii* supernatants, an effect absent in *S. cerevisiae*. This confirms previous findings that *S. boulardii* functions as an AhR agonist (27), and establishes AhR activation as a conserved feature across all tested *S. boulardii* strains. These results suggest the presence of strain-specific secreted AhR ligands. One candidate class includes tryptophan-derived indole metabolites, which are known to regulate gut immune homeostasis via AhR signalling (49). Notably, lineage-specific variants in *BNA2* and *ARO8*, genes involved in tryptophan metabolism, may contribute to the production of such ligands, providing a genomic basis for the immunomodulatory activity of S*. boulardii*. Quantitative profiling of these metabolites and functional validation in AhR reporter assays may help delineate alternative host interaction pathways, extending probiotic mechanisms beyond short-chain fatty acid production.

Consistent with these immunoregulatory features, only *S. boulardii* enhanced epithelial barrier integrity in Caco-2 monolayers, while *S. cerevisiae* had no measurable effect. Both acetate and AhR activation have been independently shown to strengthen tight junctions and reinforce barrier function (50, 51), implicating these pathways as potential mediators of the barrier-enhancing effect of *S. boulardii*.

Overall, this study demonstrates that *S. boulardii* strains from different sources are largely genomically and functionally similar, supporting their general interchangeability. Shared traits, including stress tolerance, acetate production, and anti-inflammatory activity, reinforce their probiotic potential. However, despite high genomic similarity, variability in performance may arise due to differences in manufacturing or culturing conditions (52–54), as suggested by divergent codon usage. In contrast, clear functional distinctions between *S. boulardii* and *S. cerevisiae*, particularly in stress response and immune modulation, highlight important species-specific traits. Given the increasing interest in yeast-based probiotics and advanced microbiome therapeutics (55, 56), regulatory frameworks should account for both species- and strain-level functional traits. Ensuring consistency in genomic and phenotypic profiles will be critical for quality control, efficacy, and safety across commercial formulations. While this study integrates multi-dimensional comparisons, *in vivo* validation of strain-specific effects and metabolite profiles will be essential to confirm their physiological relevance and therapeutic potential.

## Acknowledgement

This work received funding from the Novo Nordisk Foundation under NNF Grant No. NNF20CC0035580, the NNF Challenge Programme CAMiT under Grant No. NNF17CO0028232 and the Distinguished Innovator Grant No. NNF220C0081058. The authors thank Perseus for early discussions on sequencing analysis and for assistance with DNA extraction. We are also grateful to the DTU Biosustain Analytics Core for support with HPLC analysis, to Adam Jenofalvi for valuable feedback on the AhR assay, and to Ruben Vazquez-Uribe, Lise Ryberg, and Ejner Jensen for their contributions to the initial project discussions.

## Author contributions

**Hannah Elena Duffey**: Data curation; Formal analysis; Investigation; Visualisation; Methodology; Writing—original draft; Writing—review and editing. **Karl Alex Hedin**: Conceptualisation; Funding acquisition; Project administration and supervision; Data curation; Formal analysis; Investigation; Visualisation; Methodology; Writing—original draft; Writing—review and editing. **Hitesh P. Gelli**: Conceptualisation; Data curation; Investigation; Methodology. Writing—review and editing. **Troels Holger Vaaben**: Conceptualisation; Data curation; Investigation; Visualisation; Methodology; Formal analysis; Writing—review and editing. **Morten Otto Alexander Sommer**: Funding acquisition; Project administration and supervision; Writing—review and editing.

## Data availability statement

The whole genome sequencing for this study has been deposited in the National Center for Biotechnology Information (NCBI) under accession number PRJNA1312645 (https://www.ncbi.nlm.nih.gov/sra/PRJNA1312645). The remaining data are available within the article, supplementary information, or upon reasonable request from the correspondence.

## Declaration of generative AI and AI-assisted technologies

During the preparation of this manuscript, the authors used ChatGPT-4 and ChatGPT-5 to improve readability and refine the language in certain sections. All content generated with these tools was subsequently reviewed and edited by the authors, who take full responsibility for the final published version.

